# Broadly effective ACE2 decoy proteins protect mice from lethal SARS-CoV-2 infection

**DOI:** 10.1101/2023.02.22.529625

**Authors:** Mengjia Lu, Weitong Yao, Yujun Li, Danting Ma, Zhaoyong Zhang, Haimin Wang, Xiaojuan Tang, Yanqun Wang, Chao Li, Dechun Cheng, Hua Lin, Yandong Yin, Jincun Zhao, Guocai Zhong

## Abstract

As SARS-CoV-2 variants have been causing increasingly serious drug resistance problem, development of broadly effective and hard-to-escape anti-SARS-CoV-2 agents is in urgent need. Here we describe further development and characterization of two SARS-CoV-2 receptor decoy proteins, ACE2-Ig-95 and ACE2-Ig-105/106. We found that both proteins had potent and robust *in vitro* neutralization activities against diverse SARS-CoV-2 variants including Omicron, with an average IC_50_ of up to 37 pM. In a stringent lethal SARS-CoV-2 infection mouse model, both proteins lowered lung viral load by up to ∼1000 fold, prevented the emergence of clinical signs in >75% animals, and increased animal survival rate from 0% (untreated) to >87.5% (treated). These results demonstrate that both proteins are good drug candidates for protecting animals from severe COVID-19. In a head-to-head comparison of these two proteins with five previously-described ACE2-Ig constructs, we found that two of these constructs, each carrying five surface mutations in the ACE2 region, had partial loss of neutralization potency against three SARS-CoV-2 variants. These data suggest that extensively mutating ACE2 residues near the RBD-binding interface should be avoided or performed with extra caution. Further, we found that both ACE2-Ig-95 and ACE2-Ig-105/106 could be produced to gram/liter level, demonstrating the developability of them as biologic drug candidates. Stress-condition stability test of them further suggests that more studies are required in the future to improve the stability of these proteins. These studies provide useful insight into critical factors for engineering and preclinical development of ACE2 decoys as broadly effective therapeutics against diverse ACE2-utilizing coronaviruses.

**Abstract Importance:** Engineering soluble ACE2 proteins that function as a receptor decoy to block SARS-CoV-2 infection is a very attractive approach to broadly effective and hard-to-escape anti-SARS-CoV-2 agents. This study here describes development of two antibody-like soluble ACE2 proteins that broadly block diverse SARS-CoV-2 variants including Omicron. In a stringent COVID-19 mouse model, both proteins successfully protected >87.5% animals from lethal SARS-CoV-2 infection. In addition, a head-to-head comparison of the two constructs developed in this study with five previously-described ACE2 decoy constructs were performed here. Two previously-described constructs with relatively more ACE2-surface mutations were found with less robust neutralization activities against diverse SARS-CoV-2 variants. Further, the developability of the two proteins as biologic drug candidates was also assessed here. This study provides two broadly anti-SARS-CoV-2 drug candidates and useful insight into critical factors for engineering and preclinical development of ACE2 decoy as broadly effective therapeutics against diverse ACE2-utilizing coronaviruses.

**Tweet:** Two antibody-like ACE2 decoy proteins could block diverse SARS-CoV-2 variants and prevent animals from severe COVID-19.

## Introduction

The coronavirus disease 2019 (COVID-19) pandemic, which is caused by the severe acute respiratory syndrome coronavirus 2 (SARS-CoV-2), has triggered unprecedentedly rapid development of a number of countermeasures against COVID-19, including multiple prophylactic vaccines and a number of convalescent patient-derived monoclonal antibodies in clinical use^1,2^. In spite of these great achievements, SARS-CoV-2 has caused more than 600 million confirmed infections and over 6.5 million documented deaths, and the pandemic is still ongoing. This is because of the continuous emergence of new SARS-CoV-2 variants. Since late 2020, five rapid spreading and immune evasive variants of concern (VOCs; Alpha, Beta, Gamma, Delta, and Omicron) have sequentially caused multiple waves of global transmission and infections, because each of these major VOCs has more and more amino acid substitutions that have affected transmissibility and sensitivity to infection- or vaccine-induced neutralizing antibodies^3^.

SARS-CoV-2 utilizes ACE2 as a key cellular receptor to infect cells^4^. The receptor binding domain (RBD) of the viral Spike protein is responsible for the interaction and binds ACE2 with high affinity^5^. Antibodies targeting the interactions between ACE2 and SARS-CoV-2 Spike receptor-binding domain (RBD) efficiently neutralize SARS-CoV-2 infection and reduce viral load in animal models and COVID-19 patients^6-10^. So far, there are twelve anti-SARS-CoV-2 monoclonal antibodies and four two-antibody cocktails approved for clinical use^11,12^. These antibody therapeutics offer a treatment option for individuals with severe COVID-19 and are especially important for high-risk individuals where vaccination is not very effective. However, the continuous emergence of SARS-CoV-2 variants with more and more mutations in the Spike RBD region has been causing increasingly serious drug resistance issues. The original Omicron variant BA.1, first detected in November 2021, has been found with great resistance to the majority of the approved anti-SARS-CoV-2 antibody therapeutics, including ten monoclonal antibodies and three antibody cocktails^12^. Then monoclonal antibodies capable of neutralizing the original Omicron variant now have been found largely inactive against the latest new Omicron BQ and XBB subvariants, which are currently causing most new infections^12,13^. This makes the antibody therapeutics once very useful for high-risk (e.g. the elderly and immunocompromised) individuals not a good option for these individuals now. In addition, SARS-CoV-2 has been found easy to develop resistance to small molecule inhibitors such as remdesivir and nirmatrelvir^14,15^. Therefore, the development of broadly effective and hard-to-escape anti-SARS-CoV-2 agents is in urgent need.

Receptor decoy is a very promising strategy toward broadly antiviral therapeutics and has been previously applied to the development of very potent, exceptionally broad, and difficult-to-escape HIV-1 entry inhibitors^16-18^. ACE2-Ig, a recombinant Fc fusion protein of soluble human ACE2, could function as a decoy to compete with cell surface ACE2 receptor and thus should broadly block entry of diverse SARS-CoV-2 variants and difficult to be escaped. We previously described the development of improved ACE2-Ig proteins that potently neutralized the prototype SARS-CoV-2 *in vitro*^19,20^. We also demonstrated in an adenovirus-hACE2-sensitized mouse model that one of our early ACE2-Ig constructs is both prophylactically and therapeutically active against SARS-CoV-2 infection *in vivo*^21^. Here, we further optimized our ACE2-Ig constructs and assessed *in vivo* efficacy of two optimized proteins against lethal SARS-CoV-2 infection in a stringent K18-hACE2 mouse model^22^.

## Results

### ACE2-Ig-95 and ACE2-Ig-105/106 showed robust and potent *in vitro* neutralization potency against pseudoviruses of diverse SARS-CoV-2 variants

We previously described multiple ACE2-Ig constructs, ACE2-Ig-v0, ACE2-Ig-v1, ACE2-Ig-v1.1, and ACE2-Ig-v3 (**Fig 1A**). ACE2-Ig-v0 is a homo-dimeric human ACE2 peptidase domain (aa 18-615) Fc-fusion protein. ACE2-Ig-v1 carries both the peptidase domain and the Collectrin-like domain (CLD; aa 616-740) of ACE2. ACE2-Ig-v1.1 is an ACE2 D30E mutant of ACE2-Ig-v1. ACE2-Ig-v3 is an antibody-like fusion protein wherein the Fv portions of both the heavy and light chains of human IgG1 have been replaced with the ACE2 portion of the ACE2-Ig-v1.1 construct. Here we slightly modified ACE2-Ig-v1.1 by removing three amino acids (Gly-Pro-Glu) encoded by a non-self BspEI restriction site between the ACE2 domain and the IgG1 hinge, and then introducing a C-to-S mutation at position 5 of the hinge region to give the hinge more flexibility. The resulted new construct was named as ACE2-Ig-95 (**Fig 1A**). Then because residues 725-740 of ACE2 ectodomain is unstructured and might be able to serve as a linker^23^, we modified ACE2-Ig-v3 by gradually shortening the non-self (GGGGS)×3 linker between ACE2 and the kappa light chain constant domain (CL), as well as between ACE2 and the first constant domain of the IgG1 heavy chain (CH1). Compared to ACE2-Ig-v3, a new construct that has the non-self (GGGGS)×3 linker completely removed showed similar neutralization potency but improved *in vivo* pharmacokinetics profiles, including increased initial plasma concentration and extended plasma half-life (**Fig S1**). This new antibody-like construct was named as ACE2-Ig-105/106 and kept for further analysis (**Fig 1A**). In an *in vitro* pseudovirus neutralization assay using a prototype SARS-CoV-2 (WHU01) pseudovirus, similar to what we observed in the previous study^20^, ACE2-Ig-105/106 showed about 100-fold improvement over ACE2-Ig-v0 and about 10-fold improvement over ACE2-Ig-95 (**Fig 1B**). Similar trend was reproduced with the pseudoviruses of the SARS-CoV-1 and a SARS-CoV-2-like coronavirus of pangolin origin (**Fig S2**).

**Fig. 1.**
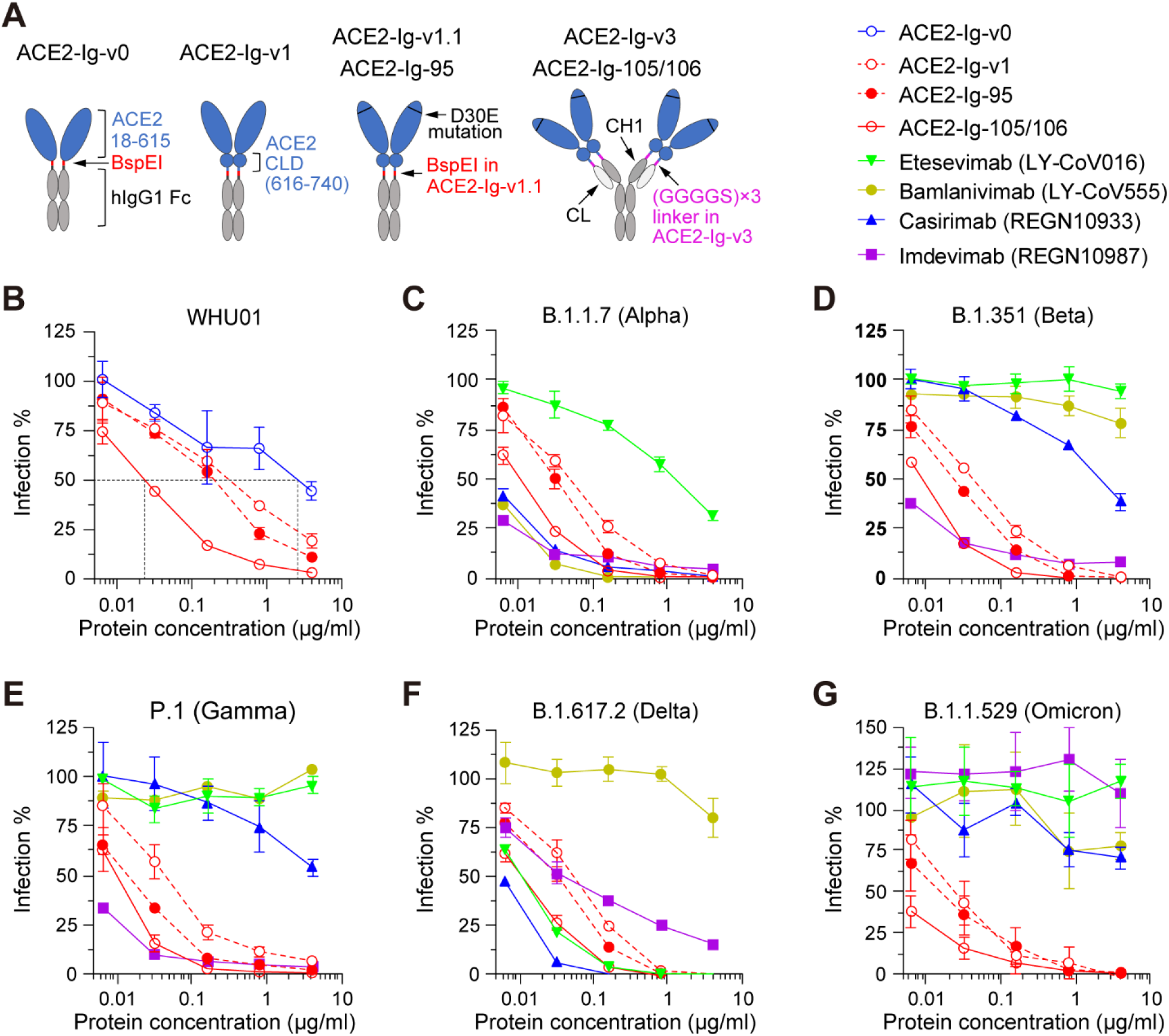
ACE2-Ig-95 and -105/106 proteins but not monoclonal antibodies robustly neutralized pseudoviruses of diverse SARS-CoV-2 variants of concern. (**A**) Diagrams showing recombinant ACE2-Ig constructs characterized in the following studies. The non-self BepEI restriction site encoding three amino acids (Gly-Pro-Glu) was removed in ACE2-Ig-95, which also has a C-to-S mutation at position 5 of the hinge region. The non-self (GGGGS)×3 linker was completely removed in ACE2-Ig-105/106. CLD, Collectrin-like domain; CH1, human IgG1 antibody heavy-chain constant domain 1; CL, human antibody kappa light-chain constant domain. (**B-G**) The indicated ACE2-Ig constructs were compared with four previously approved anti-SARS-CoV-2 monoclonal antibodies for their *in vitro* neutralization potencies against pseudoviruses of six SARS-CoV-2 variants in HeLa-hACE2 cells, a stable cell line that overexpresses human ACE2. Pseudovirus infection-mediated luciferase reporter expression was measured at 48 hours post-infection. Luciferase signals observed at each inhibitor concentration were divided by the signals observed at concentration zero to calculate percentage-of-infection (Infection %) values. Data shown are representative of three independent experiments performed by two different people with similar results, and data points represent mean ± s.d. of three biological replicates.

We then moved forward with ACE2-Ig-v1, ACE2-Ig-95, and ACE2-Ig-105/106, and compared them with four previously approved anti-SARS-CoV-2 monoclonal antibodies (etesevimab/LY-CoV016, bamlanivimab/LY-CoV555, casirivimab/REGN10933, and imdevimab/REGN10987)^6-10^ for their *in vitro* neutralization potencies against pseudoviruses of diverse SARS-CoV-2 variants. Luciferase reporter viruses pseudotyped with one of fifteen SARS-CoV-2 Spike variants, including that of VOCs Alpha (B.1.1.7), Beta (B.1.351), Gamma (P.1), Delta (B.1.617.2), and Omicron (B.1.1.529), were tested here. We found that, while most of the RBD-mutated variants showed strong or complete resistance to at least one antibody and the B.1.1.529 variant showed strong resistance to all the four tested antibodies, all the variants showed strong neutralization sensitivity to the three tested ACE2-Ig proteins (**Fig 1C-G and S3**). More importantly, compared to the prototype WHU01 variant, most tested variants showed increased neutralization sensitivity rather than any resistance to all three ACE2-Ig constructs (**Fig 1 and S3**). This might be explained by increased affinity of SARS-CoV-2 variants to human ACE2 or increased accessibility of the RBDs of SARS-CoV-2 variants^5,13,24-26^. These data demonstrate that our ACE2-Ig constructs are good drug candidates against diverse SARS-CoV-2 variants that emerged over the course of the pandemic.

### ACE2-Ig proteins with more ACE2 surface mutations neutralized SARS-CoV-2 variants less robustly

Introducing surface mutations to enhance ACE2-RBD interaction is a commonly used approach to engineer improved ACE2 decoy against SARS-CoV-2^19,20,27-30^. However, we then hypothesized that extensive ACE2 surface mutations might cause loss of neutralization potency when the heavily mutated Spike variants emerge and, more importantly, may increase the chance of eliciting anti-drug antibody (ADA) immune responses when used *in vivo*, we therefore intentionally gave up this approach at a very early stage and instead improved the proteins neutralization potency by leveraging the avidity effect of antibody-like configurations. Here, we did a head-to-head comparison of our ACE2-Ig constructs with multiple surface-mutated dimeric soluble ACE2 constructs, including one from Chan *et al*^27^, named here as ACE2-Ig-Chan-v2.4, and four from Glasgow *et al*^28^, named here as ACE2-Ig-Glasgow-293, ACE2-Ig-Glasgow-310, ACE2-Ig-Glasgow-311, and ACE2-Ig-Glasgow-313. Each of these surface-mutated ACE2-Ig constructs carries three to five ACE2 mutations (**Fig S4**) designed to enhance ACE2 interaction with the Spike protein of the prototype SARS-CoV-2 variant^27,28^.

Here, these ACE2-Ig constructs were tested for their *in vitro* neutralization potency against sixteen SARS-CoV-2 pseudoviruses, each carried a SARS-CoV-2 Spike mutant (**Fig 2 and S5**). We got multiple interesting findings here. First, although most of the surface-mutated ACE2-Ig constructs except for ACE2-Ig-Glasgow-293 showed neutralization potencies similar to that of ACE2-Ig-105/106 in most cases (**Fig 2A-E and S5**), ACE2-Ig-Glasgow-310 and ACE2-Ig-Glasgow-313 each showed significantly weaker neutralization potency against two variants (**Fig 2F-H**). More importantly, compared to the prototype variant WHU01, the mink-associated variant Y453F-F486L-N501T seemed to have partial resistance to ACE2-Ig-Glasgow-310 and ACE2-Ig-Glasgow-313 (**Fig 2H**). Note that, in contrast to ACE2-Ig-105/106 which has only one very mild D30E mutation in the ACE2 region, ACE2-Ig-Glasgow-310 and ACE2-Ig-Glasgow-313 have five ACE2 surface mutations (**Fig S4**). These data are clear evidence supporting our initial intention of avoiding intensive ACE2 surface mutations. Second, when the IC_50_ values from studies of Figs 1, 2, S3, and S5 are analyzed for each ACE2-Ig protein and antibody, ACE2-Ig-105/106 (average IC_50_ = 37 pM), ACE2-Ig-Chan-v2.4 (average IC_50_ = 59 pM), and ACE2-Ig-Glasgow-311 (average IC_50_ = 58 pM) showed the best neutralization potencies and most concentrated IC_50_ distributions against diverse SARS-CoV-2 variants (**Fig 2I**). The more scattered IC_50_ distribution observed with ACE2-Ig-Glasgow-293 and ACE2-Ig-Glasgow-313 again support our initial intention of avoiding intensive ACE2 surface mutations. These data suggest that our ACE2-Ig constructs are more likely to maintain neutralization potency against new SARS-CoV-2 variants that emerge in the future.

**Fig. 2.**
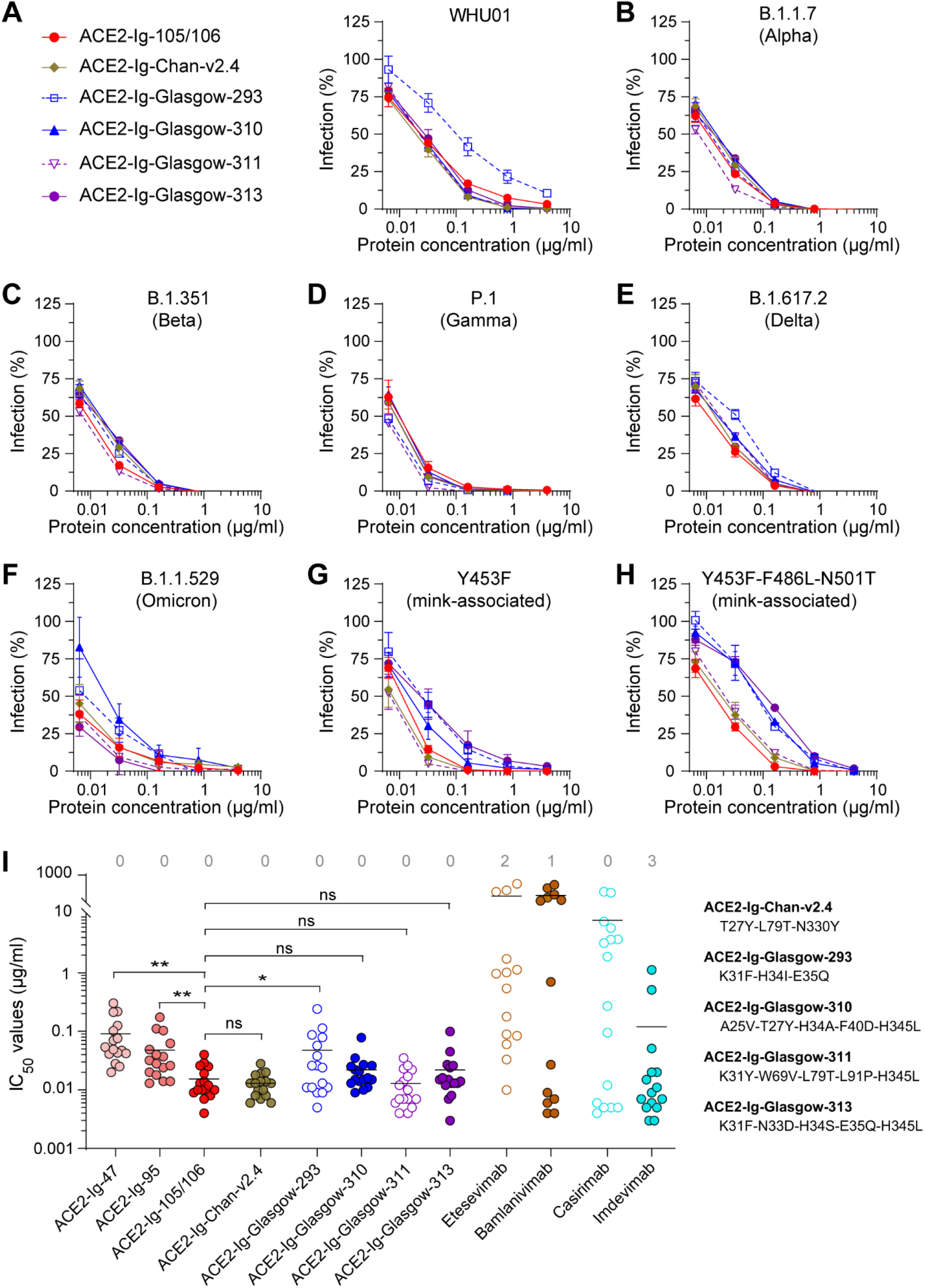
A head-to-head comparison of ACE2-Ig-105/106 with five previously published ACE2-Ig constructs. (**A-H**) Pseudovirus neutralization experiments similar to Figs 1B-G were performed to evaluate the neutralization potency and robustness of ACE2-Ig-105/106 and five surface-mutated dimeric soluble ACE2 constructs, including one from Chan *et al*^*27*^, named here as ACE2-Ig-Chan-v2.4, and four from Glasgow *et al*^28^, named here as ACE2-Ig-Glasgow-293, -310, -311, and -313. Data shown are representative of three independent experiments performed by two different people with similar results, and data points represent mean ± s.d. of three biological replicates. (**I**) The IC50 values from studies of Figs 1B-G, 2A-H, S3 and S5 are plotted. Each dot represents a SARS-CoV-2 variant. The numbers of SARS-CoV-2 variants resistant to 500 μg/mL of the indicated inhibitors are indicated at the top. Geometric means are calculated for neutralized isolates and indicated with horizontal lines. The ACE2 mutations of the five surface-mutated ACE2-Ig constructs are shown to the right of the figure. Two-sample *t*-tests (one-sided) were performed for the indicated groups, and statistical significance was indicated (ns, no significance; *, *P*<0.05; **, *P*<0.01).

### ACE2-Ig-105/106 administered intranasally most efficiently lowed lung viral load in an Ad5-hACE2-sensitized COVID-19 mouse model

We then generated two stable CHO cell pools that express ACE2-Ig-95 and ACE2-Ig-105/106, respectively. In a three-liter scale-up culture experiment, both cell pools grew well with high cell viability during a 14-day culture period (**Fig 3A and S6A**). The yield of ACE2-Ig-95 and ACE2-Ig-105/106 reached 1.6 g/L and 0.4 g/L, respectively (**Fig 3B**). Because the isoelectric points (pI) of ACE2-Ig-95 and ACE2-Ig-105/106 are 5.65 and 5.62, respectively, purified proteins were then prepared in three different buffer formulations (F1, F2, and F3) and tested for stability under three different stress conditions (free-thaw, shear flow, and temperature). Although both proteins were found sensitive to temperature stress, ACE2-Ig-105/106 showed better stability than ACE2-Ig-95 (**Fig S6B-E**). Because both proteins behaved best in buffer F3 (40 mg/mL trehalose, 0.2 mg/mL polysorbate 80, 10 mM Tris-HCl, pH7.5) under all three stress conditions nonetheless, ACE2-Ig-95 and ACE2-Ig-105/106 were then prepared in buffer F3 for animal studies.

**Fig. 3.**
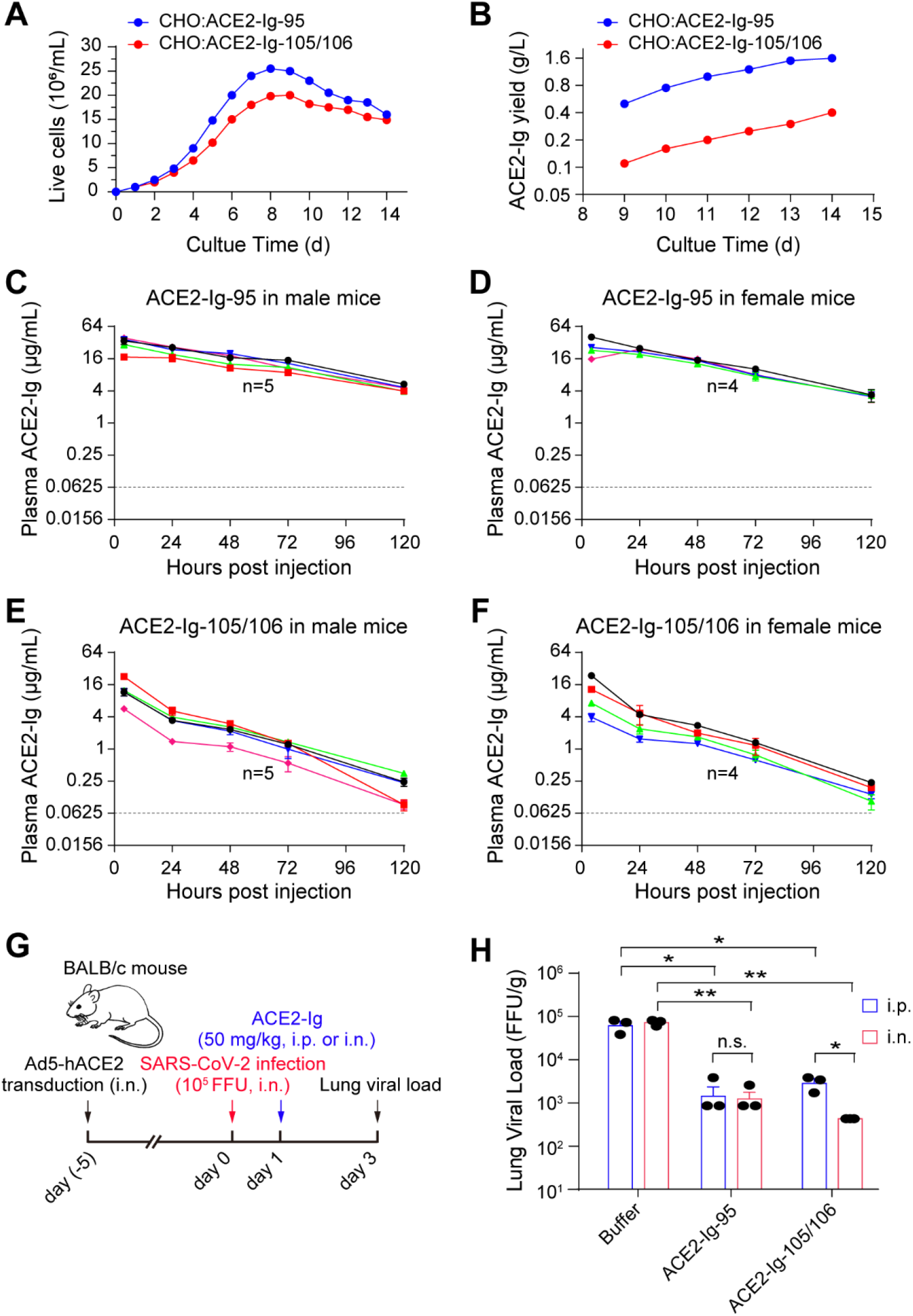
Large-scale production of ACE2-Ig-95 and -105/106 and pilot experiments exploring administration routes of the proteins for *in vivo* efficacy studies. (**A-B**) Two stable CHO cell pools that express ACE2-Ig-95 and ACE2-Ig-105/106, respectively, were generated and tested in a three-liter scale-up culture experiment. Live cell density (A) and protein yield (B) were monitored at the indicated time points. (**C-F**) Male and female BALB/c mice were injected intraperitoneally (i.p.) with 14 mg/kg of ACE2-Ig-95 or ACE2-Ig-105/106 protein for protein half-life measurement. Blood samples were collected at the indicated time points and quantitative ELISA was performed to detect the corresponding ACE2-Ig proteins from plasma samples. Each solid line represents an animal. The dash lines represent the lower limit of detection of the quantitative ELISA assay. (**G-H**) ACE2-Ig-95 and ACE2-Ig-105/106 were tested in an Ad5-hACE2-sensitized COVID-19 mouse model^33^ and administration routes (i.p. vs i.n.) for the proteins were compared. On day 1 post SARS-CoV-2 infection, animals were i.p. or i.n. treated with ACE2-Ig-95 or ACE2-Ig-105/106 at 50 kg/kg. Mice were then sacrificed on day 3 post infection and SARS-CoV-2 viral load in the lung tissue was measured using a focus forming assay. Data in G are presented as mean ± s.d. of the lung viral load data from three animals per group. Two-sample *t*-tests (one-sided) were performed for the indicated groups, and statistical significance was indicated (ns, no significance; *, *P*<0.05; **, *P*<0.01).

We first measured plasma half-life of the two proteins in mice. Both male and female BALB/c mice were injected intraperitoneally with 14 mg/kg of ACE2-Ig-95 or ACE2-Ig-105/106 protein. Blood samples were collected periodically. Quantitative ELISA detection of the corresponding ACE2-Ig proteins from plasma samples showed that ACE2-Ig-95 and ACE2-Ig-105/106, respectively, have half-lives of 43.0 ± 1.8 hours and 20.4 ± 0.3 hours (**Fig 3C-F**), both are markedly longer than the half-life of recombinant ACE2 proteins without an Fc fusion^31,32^. Because ACE2-Ig-105/106 has better *in vitro* neutralization potency but shorter plasma half-life than ACE2-Ig-95, we then performed a pilot experiment using an Ad5-hACE2-sensitized COVID-19 mouse model to compare intraperitoneal (i.p.) administration with intranasal (i.n.) administration of the two proteins for the treatment of SARS-CoV-2 infection^33^ (**Fig 3G**). Each ACE2-Ig protein at 50 mg/kg was administered intraperitoneally or intranasally to three mice per group on day 1 post SARS-CoV-2 infection. Mice were then sacrificed on day 3 post infection and the lungs were harvested for measuring viral load. Among different treatments, both i.p. and i.n. administered ACE2-Ig-95 reduced SARS-CoV-2 viral load in the lung by ∼1.5 log (**Fig 3H**). ACE2-Ig-105/106 was found significantly more effective when administered via the i.n. route, and this reduced lung viral load by ∼2 log (**Fig 3H**).

### Therapeutic use of ACE2-Ig-95 and ACE2-Ig-105/106 lowered lung viral load and improved lung histopathology in a K18-hACE2 COVID-19 mouse model

Based on the encouraging results from the pilot *in vivo* protection experiment, we decided to perform more detailed evaluation of both proteins in K18-hACE2 mouse model^22^, a more commonly used and more stringent COVID animal model. As i.n. administration was shown to be comparable or superior to i.p. administration in the pilot *in vivo* protection experiment, i.n. administration was opted for in the following studies (**Fig 4A**). Forty-eight K18-hACE2 mice were first intranasally infected with SARS-CoV-2 Hong Kong Isolate (hCoV-19/Hong Kong/VM20001061/2020) at 5000 plaque forming units (PFU). Mice were then divided into 8 groups and treatment was initiated at 6 hours post infection. Six mice per group were treated daily for five consecutive days with either buffer, etesevimab at 25 mg/kg as a positive control, or ACE2-Ig-95 or ACE2-Ig-105/106 at 4, 10, or 25 mg/kg. Mice were then sacrificed on day 5 post infection and the lungs were harvested for measuring viral load and histopathological changes. Compared to the buffer control, both proteins showed dose-dependent reduction of lung viral load and ∼3 log reduction was observed at the 10 mg/kg and 25 mg/kg doses of both proteins (**Fig 4B**). ACE2-Ig-105/106 at both 10 mg/kg and 25 mg/kg robustly lowered lung viral load in all the animals to levels close to the average lower limit of detection, indicating that the therapeutic effect of ACE2-Ig-105/106 has plateaued at the 10 mg/kg dose (**Fig 4B**). As there’s no significant difference in the viral loads between the animals treated with positive control (etesevimab at 25 mg/kg) and ACE2-Ig-95 or -105/106 (4 mg/kg or 10 mg/kg), both ACE2-Ig proteins are considered more effective than etesevimab.

**Fig. 4.**
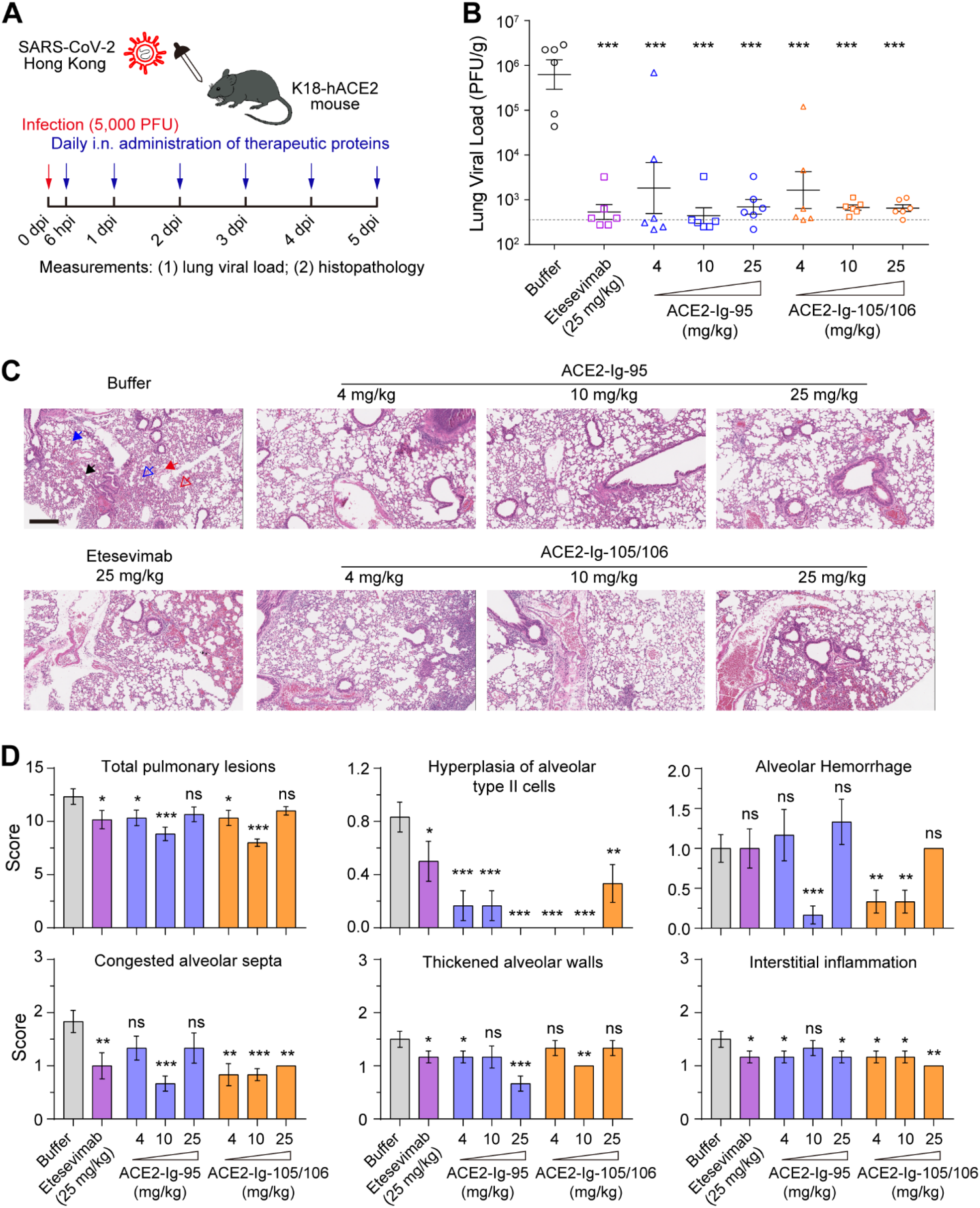
ACE2-Ig-95 and ACE2-Ig-105/106 lowered lung viral load and improved lung histopathology in SARS-CoV-2-infected K18-hACE2 mice. (**A**) A diagram representing the experimental design used in the following animal studies. (**B-D**) K18-hACE2 mice treated following the procedure in A (n=6 per group) were sacrificed on day 5 post infection and the lungs were harvested for measuring viral load (B) and histopathological changes (C-D). Each data point in B represents an animal, and data are presented as mean ± s.d. of the lung viral load data from six animals per group. The dash line represents the lower limit of detection of the lung viral load assay. Compared to the buffer control, all the treatments significantly lowered lung viral load (two-sample *t*-tests, one-sided; ***, *P*<0.001). No significance was found among treatment groups (B). The scale bar in C represents 300 μm. Pathological changes were indicated with different arrows. Blue solid arrow, hyperplasia of alveolar type II (ATII) cells; blue blank arrow, congested alveolar septa; red solid arrow, interstitial inflammation; red blank arrow, alveolar hemorrhage; black arrow, thickened alveolar walls. Data points in D represents mean ± s.e.m of the pathological scores obtained from two sections per animal, and six animals per group. Two-sample *t*-tests (one-sided) were performed between the buffer control group and each treatment group. Statistical significance was indicated (ns, no significance; *, *P*<0.05; **, *P*<0.01; ***, *P*<0.001). Pathological score data for the lesions that no significant difference was found between control and treatment groups are shown in Fig S7.

Lung histopathology analysis showed that, compared to buffer-treated mice, all the drug-treated groups showed improvement in pulmonary lesions and lower pathological scores (**Fig 4C and D**). Consistent with what we observed with the lung viral load data, mice treated with either of the ACE2-Ig proteins at 10 mg/kg showed more significant histopathological improvement than animals treated with 25 mg/kg of etesevimab, again demonstrating that both ACE2-Ig proteins are more effective than etesevimab (**Fig 4D**). When each pulmonary lesion was analyzed individually, multiple lesions including hyperplasia of alveolar type II cells, alveolar hemorrhage, congested alveolar septa, thickened alveolar walls, and interstitial inflammation were found to be significantly improved by the ACE2-Ig proteins (**Fig 4D and S7**).

### ACE2-Ig-95 and ACE2-Ig-105/106 effectively protected K18-hACE2 mice from lethal SARS-CoV-2 infection

We then performed another K18-hACE2 mouse experiment to assess the ability of ACE2-Ig-95 and ACE2-Ig-105/106 to save animals from infection-caused clinical signs and fatality. In this experiment, sixty-four SARS-CoV-2-infected K18-hACE2 mice were divided into eight treatment groups. Eight mice per group were treated daily for seven consecutive days with either buffer, etesevimab at 25 mg/kg, or ACE2-Ig-95 or ACE2-Ig-105/106 at 4, 10, or 25 mg/kg (**Fig 5A**). Mice were continuously monitored from day 0 through day 14 post infection for body weight, clinical signs of SARS-CoV-2 infection, and survival. All eight mice in the buffer control group showed marked (up to 15%) body weight loss by day 7 post infection. In contrast, the majority of the forty-eight ACE2-Ig-treated animals across the three treatment doses did not display infection-associated significant weight loss (**Fig 5B**). Differences in bodyweight loss between the buffer control group and each treatment group are all significant on days 6 post infection (two-sample *t*-tests, one-sided, *P*<0.05).

**Fig. 5.**
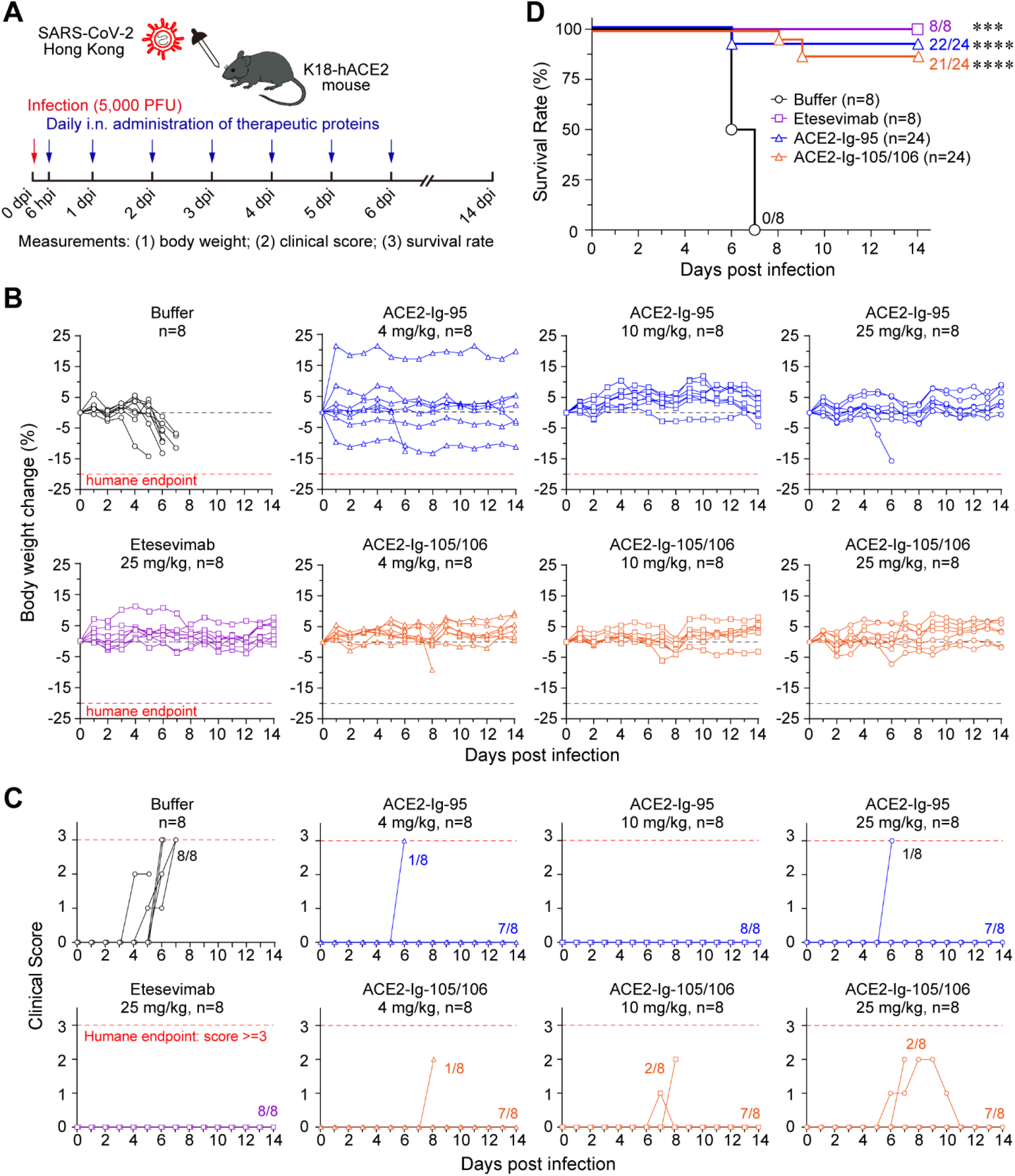
ACE2-Ig-95 and ACE2-Ig-105/106 effectively protected K18-hACE2 mice from lethal SARS-CoV-2 infection. (**A**) A diagram representing the experimental design used in the following animal studies. (**B-D**) K18-hACE2 mice treated following the procedure in A (n=8 per group) were continuously monitored from day 0 through day 14 post infection for body weight (B), clinical signs of SARS-CoV-2 infection (C), and survival (D). Each solid line in B and C represents an animal, and the red dash lines represent the human endpoints for the studies. Differences in bodyweight loss between the buffer control group and each treatment group are all significant on days 6 post infection (two-sample *t*-tests, one-sided, *P*<0.05). For clinical score analysis, clinical signs including piloerection, hunched posture, decreased activity, and respiration difficulty were monitored and scored. The number of animals displayed clinical signs in each group was indicated. Differences in clinical signs between the buffer control group and each treatment group are all significant on days 5-7 post infection (two-sample *t*-tests, one-sided, *P*<0.01). For survival analysis, data for animals treated with different doses of a same protein (ACE2-Ig-95 or ACE2-Ig-105/106) were pooled together in the Kaplan-Meier survival curves. The number of animals survived was indicated at the terminal point of each group. Log-rank (Mantel-Cox) test was performed to determine the statistical significance between the control and each treatment groups (***, *P*<0.001;****, *P*<0.0001)

For clinical score analysis, clinical signs including piloerection, hunched posture, decreased activity, and respiration difficulty were monitored and scored. The presence of each sign gave an animal a score of 1. The sum of the scores of an animal was defined as the animal’s clinical score. A clinical score of 3 or more was considered as a humane endpoint. The clinical scores for each animal were plotted against the time of monitoring (**Fig 5C**). Consistent with the body weight changes, all eight mice in the buffer control group displayed multiple SARS-CoV-2 infection-associated clinical signs and therefore high clinical scores, and most of them had clinical scores reached the human endpoint, by day 7 post infection. Treatment with ACE2-Ig-95 or ACE2-Ig-105/106 eliminated the development of clinical signs in most of the animals and differences in clinical signs between the buffer control group and each treatment group are all significant on days 5-7 post infection (two-sample *t*-tests, one-sided, *P*<0.01). Specifically, two out of twenty-four ACE2-Ig-95-treated mice displayed severe clinical signs that reached the humane endpoint on day 6 post infection. Although more (five out of twenty-four) animals treated with ACE2-Ig-105/106 displayed clinical signs, there was likely a delay in the onset of the clinical signs and the signs finally resolved in two of these five animals.

According to the IACUC protocol, animals with more than 20% weight loss or a clinical score of 3 or more would be euthanized and considered as infection-caused fatalities. In the buffer control group, two of the mice died and the other six reached clinical score humane endpoint by day 7 post infection. The median survival time of this group is 6.5 days post infection (**Fig 5D**). In contrast, only two out of twenty-four animals treated by ACE2-Ig-95, one in the 4 mg/kg group and the other in the 25 mg/kg group, reached clinical score humane endpoint on day 6 post infection. Three out of twenty-four animals treated by ACE2-Ig-105/106, one in each dose group, died on day 8 or 9 post infection (**Fig 5D and S8**). Therefore, without treatment, the survival rate of these SARS-CoV-2 infected animals was 0%. Then treatment with ACE2-Ig-95 and ACE2-Ig-105/106, respectively, dramatically increased the survival rate to 91.7% (*P*<0.0001) and 87.5% (*P*<0.0001) across the three dose groups (**Fig 5D**). For the eight mice treated with10 mg/kg of ACE2-Ig-95, the survival rate reached 100% (*P*=0.0001; **Fig S8B**). These results demonstrate that both ACE2-Ig-95 and ACE2-Ig-105/106 are good drug candidates that can be used to protect animals from severe COVID-19.

## Discussion

Developing ACE2 decoy is a very promising approach to broadly effective and hard-to-escape anti-SARS-CoV-2 agents^5,19-21^. So far, all the major SARS-CoV-2 VOCs that sequentially caused multiple waves of global transmission have been found to have ACE2-binding affinities higher than the original variant^5,13,24,25^. Structural studies have shown that Spike trimers of many SARS-CoV-2 variants have significantly higher propensity to adopt ‘RBD-up’ or open state than the D614G variant does^24,26^. These are consistent with our data that, compared to the prototype SARS-CoV-2, all the tested variants showed either comparable or increased sensitivity to our ACE2-Ig constructs (**Fig 1, 2I and S3**). These findings all suggest that ACE2-Ig is likely to be a long-term viable approach for coping with diverse circulating and emerging SARS-CoV-2 variants. In addition, a number of other coronaviruses, including SARS-CoV-1, human coronavirus NL63, and some SARS-like CoVs of bat or pangolin origin also utilize ACE2 as entry receptor^34-38^. Recently, two close relatives of MERS-CoV in bats were found also utilize ACE2 as their functional receptors^39^. Considering that coronaviruses have very broad host ranges and moderate recombination frequencies^19,40,41^, it therefore can’t rule out the possibility of more coronaviruses being found to use ACE2 as their functional receptors in the future. ACE2-Ig is therefore a broadly anti-coronavirus drug candidate that should be included in the toolbox for pandemic preparedness.

To develop a potent and safe ACE2-Ig for human use, four critical factors should be explored. The first critical but initially often neglected factor is the truncation of the transmembrane ACE2 protein for soluble expression. The extracellular region of ACE2 (residues 18-740) consists of a peptidase domain (residues 18-615) and a Collectrin-like domain (CLD; residues 615-740)^23^. At the beginning of the pandemic, several groups including us independently explored the utility of ACE2-Ig decoy as an anti-SARS-CoV-2 agent^19,42-45^. While most of these studies directly opted for ACE2-Ig constructs that carried the ACE2 peptidase domain but no CLD^42-45^, we carefully compared a panel of CLD-free and CLD-containing ACE2-Ig constructs^19^. We found that the CLD-containing ACE2-Ig constructs consistently showed ∼20-fold better neutralization potency than the CLD-free ACE2-Ig constructs^19^. Since then, our ACE2-Ig studies^20,21^ (**Fig 1B and S2**) and most of the ACE2-Ig studies from other groups included the CLD in the constructs^27-29,46^. Recently, a study reported that CLD could also dramatically extend serum half-life of their human IgG4 Fc-based ACE2-Ig constructs^46^. Interestingly, with our constructs that are based on human IgG1 rather than IgG4 Fc, we did not observe this significant difference in half-lives between CLD-free and CLD-containing constructs (data not shown), suggesting the CLD’s contribution on half-life might be IgG subclass-dependent.

The second and most frequently investigated factor is mutations to ACE2 surface residues. Introducing mutations to ACE2 surface residues is a commonly adopted approach for engineering improved ACE2-Ig decoy that has enhanced RBD recognition and SARS-CoV-2 neutralization activity^19,20,27-30^. We intentionally avoided this approach at very early stage and instead improved the proteins neutralization potency by leveraging the avidity effect of antibody-like multi-valent configurations. As a result, when we previously used *in vitro* serial viral passage to study drug-induced viral escape, no ACE2-Ig escape mutant was detected in wild-type ACE2-Ig treated samples^21^. In addition, none of the SARS-CoV-2 variants in the current study showed significantly decreased sensitivity to our ACE2-Ig constructs (**Fig 1, 2 and S3**). In contrast, we observed in three SARS-CoV-2 variants seemingly partial resistance to two previously published ACE2-Ig constructs, each carrying five surface mutations in the ACE2 region (**Fig 2F-H**). These data clearly suggest that extensively mutating the ACE2 residues near the RBD-binding interface should be avoided or performed with extra caution. It might result in compromised neutralization breadth, not to mention the risk of eliciting ADA immune responses which might also target endogenous ACE2 protein.

The other two important factors are ACE2 peptidase activity and Fc effector functions. Whether the peptidase activity should be retained in an ACE2-Ig product is still under debate. Although some previous non-COVID studies in animals as well as in humans^47-49^ and our recent ACE2-Ig study in mice^21^ all support a potentially beneficial and non-antiviral role of ACE2 peptidase activity in COVID-19 treatment, more detailed *in vivo* studies to dissect the contribution of ACE2 peptidase activity to the treatment is needed in the future. Fc effector functions are normally beneficial for antibody-like antiviral biologics^50^. Indeed, a recent study reported that Fc effector functions could enhance ACE2-Ig’s therapeutic activity in a COVID-19 mouse model^29^. However, because human ACE2 is an endogenous protein with broad substrate specificity^51,52^, keeping or enhancing ACE2-Ig’s Fc effector functions might cause off-target cell killing. Therefore, more carefully designed studies to investigate this aspect should be warranted in the future. An ACE2-Ig product that keeps Fc effect functions has moved into clinical stage^53^ (NCT05116865, NCT05659602). Safety data from these trials should be informative.

In terms of the four critical factors, both ACE2-Ig-95 and ACE2-Ig-105/106 carry a CLD-containing, barely mutated, and peptidase-active ACE2 ectodomain and an effector function-competent IgG1 Fc. They neutralized diverse SARS-CoV-2 variant pseudoviruses with robust potencies (**Fig 1 and 2**) and markedly lowered lung viral load in two different COVID-19 mouse models (**Fig 3 and 4**). In the more stringent K18-hACE2 mouse model, both proteins at 4 mg/kg daily doses effectively prevented the emergence of clinical signs and greatly increased survival rate of the animals (**Fig 5**). These data demonstrate that ACE2-Ig-95 and ACE2-Ig-105/106 are broadly effective and hard-to-escape anti-SARS-CoV-2 drug candidates that can be used to protect animals from severe COVID-19. Besides efficacy, producibility and stability are critical determinants for the developability of a biologic drug candidate. Three-liter scale-up culture tests showed that both ACE2-Ig-95 and ACE2-Ig-105/106 stable cells had a yield at g/L level (**Fig 3B**), demonstrating that both proteins as biologic drug candidates are developable. However, stability tests showed that a significant fraction of both proteins, especially ACE2-Ig-95, aggregated under a temperature stress condition (**Fig S6**). Therefore, more in-depth formulation or engineering studies should be performed in the future to address the stability issue of these proteins.

## Materials and Methods

### Cells

293T cells and HeLa cells were kindly provided by Stem Cell Bank, Chinese Academy of Sciences, confirmed mycoplasma-free by the provider, and maintained in Dulbecco’s Modified Eagle Medium (DMEM, Life Technologies) at 37 °C in a 5% CO2-humidified incubator. Growth medium was supplemented with 2 mM Glutamax-I (Gibco, Cat. No. 35050061), 100 μM non-essential amino acids (Gibco, Cat. No. 11140050), 100 U/mL penicillin and 100 μg/mL streptomycin (Gibco, Cat. No. 15140122), and 10% heat-inactivated FBS (Gibco, Cat. No. 10099141C). HeLa-based stable cells expressing human ACE2 were maintained under the same culture condition as HeLa, except that 3 μg/mL of puromycin was added to the growth medium. 293F cells for recombinant protein production were generously provided by Dr. Yu J. Cao (School of Chemical Biology and Biotechnology, Peking University Shenzhen Graduate School) and maintained in SMM 293-TII serum-free medium (Sino Biological, Cat. No. M293TII) at 37 °C, 8% CO2, in a shaker incubator at 125rpm.

### Plasmids

DNA fragment encoding spike protein of SARS-CoV-2 WHU01 (GenBank: MN988668.1) was synthesized by the Beijing Genomic Institute (BGI, China) and then cloned into pCDNA3.1(+) plasmid between EcoRI and XhoI restriction sites. Plasmids encoding SARS-CoV-2 spike variants were generated according to the in-fusion cloning protocol. To facilitate SARS-CoV-2 pseudovirus production, spike sequences for WHU01 and all the variants investigated in this study all contain a deletion (ΔPRRA) or GSAS substitution at the PRRA furin-cleavage site. Our previous study showed that the ΔPRRA mutation does not affect SARS-CoV-2 cross-species receptor usage or neutralization sensitivity^20^. The retroviral reporter plasmids encoding a Gaussia luciferase reporter gene were constructed by cloning the reporter genes into the pQCXIP plasmid (Clontech). Plasmids encoding soluble ACE2 variants fused with human IgG1 Fc were described in our previous study^20^. DNA fragments encoding heavy and light chains of anti-SARS-CoV-2 antibodies were synthesized by Sangon Biotech (Shanghai, China) and then cloned into a pCAGGS plasmid.

### Production and Purification of ACE2-Ig protein and SARS-CoV-2 antibodies by transient transfection

293F cells at 6 × 10^5^ cells/mL density were seeded into 100 mL SMM 293-TII serum-free medium (Sino Biological, Cat. No. M293TII) one day before transfection. Cells were then transfected with 100 μg plasmid in complex with 250 μg PEI MAX 4000 (Polysciences, Cat. No. 24765-1). Cell culture supernatants were collected at 48 to 72 hours post-transfection. Human IgG1 Fc-containing proteins were purified using Protein A Sepharose CL-4B (GE Healthcare, Cat. No. 17-0780-01), eluted with 0.1 M citric acid at pH 4.5 and neutralized with 1 M Tris-HCl at pH 9.0. Buffers were then exchanged to PBS, and proteins were concentrated by 30 kDa cut-off Amicon Ultra-15 Centrifugal Filter Units (Millipore, Cat. No. UFC903096).

### Production of reporter retroviruses pseudotyped with SARS-CoV-2 spike variants

MLV retroviral vector-based SARS-CoV-2 spike pseudotypes were produced according to our previous study^20^, with minor changes. In brief, 293T cells were seeded at 30% density in 150 mm dish at 12-15 hours before transfection. Cells were then transfected with 67.5 μg of polyethyleneimine (PEI) Max 40,000 (Polysciences, Inc, Cat. No. 24765-1) in complex with 3.15 μg of plasmid encoding a spike variant, 15.75 μg of plasmid encoding murine leukemia virus (MLV) Gag and Pol proteins, and 15.75 μg of a pQCXIP-based luciferase reporter plasmid. Eight hours after transfection, cell culture medium was refreshed and changed to growth medium containing 2% FBS (Gibco, Cat. No. 10099141C) and 25 mM HEPES (Gibco, Cat. No. 15630080). Cell culture supernatants were collected 36-48 hours post-transfection, spun down at 3000×g for 10 min, and filtered through 0.45 μm filter units to remove cell debris. SARS-CoV-2 spike-pseudotyped viruses were then concentrated 10 times at 2000×g using 100 kDa cut-off Amicon Ultra-15 Centrifugal Filter Units (Millipore. Cat. No. UFC910024).

### Pseudovirus Titration

Pseudovirus titer were determined using a reverse transcriptase activity assay. Reverse transcriptase-containing pseudoviral particles and recombinant reverse transcriptase standard of known concentrations (TAKARA, Cat. No. RR047A) were 10-timed diluted with nuclease-free water (Invitrogen, Cat. No. 10977015) and lysed with 2× concentrated lysis buffer (0.25% Triton X-100, 50 mM KCL, 100 mM Tris-HCl pH 7.4, 40% glycerol, 1/50 volume of RNase inhibitor; NEB, Cat. No. M0314S) at room temperature for 10 min. Reverse transcription was performed according to the manufacturer’s protocol (TAKARA, Cat. No. RR047A) using 1 µL of the lysate as reverse transcriptase and TRIzol reagent-isolated 293T total RNA as template. Reverse transcription products were then subjected to qPCR with a commercial kit (TAKARA, Cat. No. RR820Q) to amplify GAPDH (Forward primer: 5’-CCACTCCTCCACCTTTGAC-3’, Reverse primer: 5’-ACCCTGTTGCTGTAGCCA-3’) in Applied Biosystems QuantStudio 5. A standard curve was generated based on qPCR Ct values obtained with serially diluted recombinant reverse transcriptase standard.

### SARS-CoV-2 pseudovirus neutralization assay

Pseudovirus neutralization experiments were performed following our previous study^20^, with minor changes. In brief, SARS-CoV-2 spike variant pseudotyped luciferase reporter viruses equivalent to 8×10^10^ U reverse transcriptase were pre-diluted in DMEM (2% FBS, heat-inactivated) containing titrated amounts of an ACE2-Ig construct or an anti-SARS-CoV-2 antibody. Virus-inhibitor mixtures were incubated at 37°C for 30 min, then added to HeLa-hACE2 cells in 96-well plates and incubated overnight at 37°C. Virus-inhibitor-containing supernatant was then removed and changed with 150 μL of fresh DMEM (2% FBS) and incubated at 37°C. Cell culture supernatants were collected for Gaussia luciferase assay at 48 h post-infection.

### Gaussia luciferase luminescence flash assay

To measure Gaussia luciferase expression, 20 μL of cell culture supernatant of each sample and 100 μL of assay buffer containing 4 μM coelenterazine native (Biosynth Carbosynth, Cat. No. C-7001) were added to one well of a 96-well black opaque assay plate (Corning, Cat. No. 3915) and measured with Centro LB 960 microplate luminometer (Berthold Technologies) for 0.1 second/well.

### Stable CHO cells generation and 3-L scale-up production of ACE2-Ig-95 and ACE2-Ig-105/106

Two CHOZN® CHO K1-based stable cell pools stably expressing ACE2-Ig-95 and ACE2-Ig-105/106, respectively, were generated and tested for 3-L scale-up production and stress-condition stability by Canton Biologics (Guangzhou, China). In brief, CHOZN® CHO K1 cells were thawed and maintained in EXCELL CD CHO Fusion medium (Sigma, Cat. No. 14365C) containing 4 mM L-glutamine at 37°C, 5% CO2, 85% humidity, with 140-rpm agitation. Cells were then transfected with ACE2-Ig-95 or ACE2-Ig-105/106 plasmid using PEI 25K (Polysciences, Cat. No. 23966-1) and selected using methionine sulfoximine (MSX)-containing EX-CELL CD CHO Fusion Medium (Sigma, Cat. No. 14365C). Cells passed the MSX selection cycles were then subjected to a pilot production experiment. Cells were maintained in 280 mL culture for 14 days in EX-CELL Advanced CHO Fed-batch Medium (Sigma, Cat. No. 14366C) that was then added with Cell boost 7a (Hyclone, Cat. No. SH31119.01) and Cell boost 7b (Hyclone, Cat. No. SH31120.01). Cell viability, live cell density, and protein expression were monitored on daily basis from day 3 through day 14 of the culture period. Protein production-validated cells were then subjected to a scale-up production experiment in 3-L bioreactors (Applikon my-Control). Cells were initially diluted to 0.5×10^6^ cells/mL in 1.2 L EX-CELL Advanced CHO Fed-batch Medium (Sigma, Cat. No. 14366C). Starting from day 3, cells were added daily with glucose to 8 g/L, 3% volume of Cell boost 7a (Hyclone, Cat. No. SH31119.01) and 0.3% volume of Cell boost 7b (Hyclone, Cat. No. SH31120.01) until day 14. Cells were maintained at 37°C, 40% dissolved oxygen, and stirred at 300rpm/320rpm with gas flow rates of 33 mL/min and 12 mL/min (0.01 vvm). Cell viability, live cell density, and protein expression (Protein A-HPLC) were monitored on daily basis through the 14-day culture period. ACE2-Ig-95 in cell culture supernatant was first captured using MabSelect SuRe affinity column (purity: 83.73%; recovery rate: 102%) and then purified using UniHR Phenyl 30L Hydrophobic Interaction Chromatography column (purity: ∼95%; recovery rate: 32%). ACE2-Ig-105/106 in cell culture supernatant was first captured using MabSelect SuRe affinity column (purity: 85%; recovery rate: 85%) and then purified using Diamond Q mustang anion exchange column (purity: ∼94%; recovery rate: 70%).

### ACE2-Ig-95 and ACE2-Ig-105/106 stability tests under stress conditions

Because the isoelectric points (pI) of ACE2-Ig-95 and ACE2-Ig-105/106 are 5.65 and 5.62, respectively, the proteins were prepared at 10 mg/mL concentration in three different buffers (F1, F2, and F3). All three buffers contain 40 mg/mL trehalose and 0.2 mg/mL polysorbate 80. In addition, buffer F1 (pH6.5) and F2 (pH7.0) have 10 mM Histidine. Buffer F3 (pH7.5) has 10 mM Tris-HCl. Proteins in these different buffers were then assessed for their stability under the following three stress conditions: freeze-thaw stress (five cycles of freezing at -80 °C and thawing at room temperature), shear stress (agitation at 300rpm, 37 °C, for one week), and temperature stress (incubation at 40 °C for two weeks). Protein samples were then subjected to size exclusion chromatography (SEC) analysis using a TSKgel G3000SW_XL_ column to quantify the fractions of high molecular weight, main peak, and low molecular weight, respectively.

### Focus forming assay (FFA) for SARS-CoV-2 quantification

All SARS-CoV-2 live virus infection experiments were performed in a Biosafety Level 3 (BSL-3) laboratory. Vero E6 cells were seeded onto 96-well plates overnight and grown into confluent monolayers. Fifty microliters of 10-fold-diluted SARS-CoV-2 stock or supernatant of lung homogenate was added into 96-well plate and adsorbed at 37°C for 1 h with agitation every 10 min. Then the virus or supernatant of lung homogenate were removed and covered with 100 μL Minimum Essential Medium (MEM) containing 1.2% Carboxymethylcellulose (1.2% CMC). Twenty-four hours post infection, the overlay was discarded and the cell monolayer was fixed with 4% paraformaldehyde solution for 2 h at room temperature. After permeabilized with 0.2% Triton X-100 for 20 min at room temperature, the plates were sequentially stained with cross-reactive rabbit anti-SARS-CoV-N IgG (Sino Biological Inc) as the primary antibody and HRP-conjugated goat anti-rabbit IgG (H + L) (Jackson ImmunoResearch) as the secondary antibody at 37°C for 1 h. The reactions were developed with KPL TrueBlue Peroxidase substrates. The numbers of SARS-CoV-2 foci were calculated using CTL ImmunoSpot S6 Ultra reader (Cellular Technology Ltd) and titers of the virus were expressed as focus forming unit (FFU) per milliliter.

### SARS-CoV-2 live virus studies in Ad5-hACE2-sensitized mice

Ad5-hACE2-sensitized mice were used to evaluate *in vivo* efficacy of ACE2-Ig-95 and ACE2-Ig-105/106 following our previous study^33^. These experiments were conducted in BSL-3 under protocols approved by the Institutional Animal Care and Use Committee (IACUC) in the Guangzhou Customs District Technology Center. Briefly, six-week-old female BALB/c mice (∼15 g) were first intranasally transduced with 2.5×10^8^ FFU of Ad5-hACE2. Five days later, animals were then intranasally challenged with 1×10^5^ FFU of SARS-CoV-2. On day 1 post SARS-CoV-2 infection, animals were intraperitoneally injected with an ACE2-Ig protein at 50 mg/kg. Animals were then sacrificed on day 3 post SARS-CoV-2 infection and the lungs were collected in PBS and homogenized. Titers of SARS-CoV-2 in clarified supernatants were determined using FFA assay in Vero E6 cells and expressed as FFU per gram of tissue.

### SARS-CoV-2 live virus studies in K18-hACE2 mice

Six-to-eight-week-old specific pathogen-free female B6.Cg-Tg(K18-ACE2)2Prlmn/J transgenic mice, called K18-hACE2 mice hereafter, were purchased from the Jackson Laboratory. All K18-hACE2 mice experiments were performed in BSL-3 under approved IACUC protocols by a Wuxi AppTech sponsored research institution. Animals were housed in individually ventilated cages and randomly assigned to different treatment groups. Each treatment group has six or eight mice. On day 0, animals were intranasally infected with 5000 PFU SARS-CoV-2 Hong Kong Isolate (Hong Kong/VM20001061/2020; ATCC) in 50 μL volume.

For lung viral load and histopathological analysis, forty-eight SARS-CoV-2-infected K18-hACE2 mice were divided into 8 groups and treatment was initiated at 6 hours post infection. Six mice per group were treated daily for five consecutive days with either buffer, etesevimab at 25 mg/kg as a positive control, or ACE2-Ig-95 or ACE2-Ig-105/106 at 4, 10, or 25 mg/kg. Mice were then sacrificed on day 5 post infection. The left lungs of the animals were harvested and fixed in 10% formalin for histopathological analysis. The right lungs of the animals were collected, weighed, and stored in EMEM medium containing 1% FBS at -80°C for lung viral load analysis.

For survival analysis, sixty-four SARS-CoV-2-infected K18-hACE2 mice were divided into eight treatment groups and treatment was initiated at 6 hours post infection. Eight mice per group were treated daily for seven consecutive days with either buffer, etesevimab at 25 mg/kg, or ACE2-Ig-95 or ACE2-Ig-105/106 at 4, 10, or 25 mg/kg. Mice were continuously monitored from day 0 through day 14 post infection for body weight, clinical signs of SARS-CoV-2 infection, and survival. For clinical sign monitoring, piloerection, hunched posture, decreased activity, and respiration difficulty were monitored and scored. The presence of each sign gave an animal a score of 1. The sum of the scores of an animal was defined as the animal’s clinical score. According to the IACUC protocol, the humane endpoint is defined as bodyweight loss of 20% or more, clinical score of 3 or more, or the agonal state. An animal that has reached the human endpoint will be euthanized.

### Lung viral load measurement using plaque assay

Vero E6 cells were seeded into 6-well plates to a 7.5×10^5^ cell/mL density before infection. The right-lung samples collected in Eagle’s Minimum Essential Medium (EMEM) with 1% FBS were homogenized in a tissue homogenizer. Samples were then centrifuged, and the supernatant was10-fold serially diluted and used to infect Vero E6 cells for 1 hour at 37°C with shaking at 15-min intervals. Cell culture supernatant was then removed and 1% agarose in EMEM supplemented with 20% FBS was added to the cells and incubated for 3 days at 37°C. Agarose was then carefully removed and cells were first fixed with 95% ethanol for 15 min. After a brief wash with PBS, cells were then fixed and stained for 15min in 10% formalin containing 1% crystal violet. SARS-CoV-2 infection-caused plaques were counted and viral titers were finally calculated and converted as plaque forming unit (PFU) per gram lung tissue.

### Lung histopathological analysis

The left lungs fixed in 10% formalin were paraffin-embedded and sectioned. Sections were stained with hematoxylin/eosin for histopathological analysis. Photomicrographs taken by Leica Aperio AT2 digital pathology scanning system (LeicaBiosystems, INC) were subjected to semi-quantitative histopathological analysis. The photomicrographs were analyzed for the presence of the following alveoli-region lesions (pulmonary edema, alveolar hemorrhage, thickened alveolar walls, alveolar inflammation, necrosis, hyaline membrane, thrombus, hyperplasia of alveolar type II cells, and alveolar space protein fragments) and mesenchyme-region lesions (interstitial inflammation, congested alveolar septa, perivascular edema, and perivascular hemorrhage). The presence of each lesion gave a photomicrograph a grade score between 0 to 5, according to the severity of the lesion. A score of 0 means no lesion, 1 means minimal, 2 means slight, 3 means moderate, 4 means marked, and 5 means severe.

### Data collection and analysis

MikroWin 2000 Software (Berthold Technologies) was used to collect luciferase assay data. Leica Aperio AT2 digital pathology scanning system (LeicaBiosystems, INC) was used to collect the photomicrographs for the lung histopathological analysis. GraphPad Prism 9.4 software was used for figure preparation and statistical analyses.

### Statistical Analysis

All the *in vitro* experiments were independently performed two or three times and data are expressed as mean values ± s.d. or s.e.m. Statistical analyses were performed using two-sample *t*-test (IC50, lung viral load and histopathology) or log-rank (Mantel-Cox) test (survival) when applicable. Differences were considered significant at *P* < 0.05. The values for n, *P*, and the specific statistical test performed for each experiment are included in the figure legend and main text.

## Author contributions

Conceptualization: GZ

Methodology: WY, YL, HW, XT, YW, CL, DC, HL, GZ

Investigation: ML, WY, DM, ZZ, YL Supervision: YL, YY, JZ, GZ Writing—original draft: GZ, ML, HW

Writing—review & editing: ML, WY, YL, DM, ZZ, HW, XT, YW, CL, DC, HL, YY, JZ, GZ

## Data and materials availability

All data are available in the main text or the supplementary materials. This study did not generate unique datasets or code. Our in-house research resources, including methods, plasmids, and protocols, are available upon reasonable request to qualified academic investigators for noncommercial research purposes. All reagents in-house developed, including vector plasmids and detailed methods, will be made available upon written request.

